# Continual familiarity decoding from recurrent connections in spiking networks

**DOI:** 10.1101/2025.01.13.632765

**Authors:** Viktoria Zemliak, Gordon Pipa, Pascal Nieters

**Affiliations:** Institute of Cognitive Science, University of Osnabrück, Osnabrück, Germany; Frankfurt Institute of Advanced Studies, Frankfurt, Germany

**Keywords:** Spiking networks, STDP, synaptic plasticity, continual familiarity, spike synchrony

## Abstract

Familiarity memory enables recognition of previously encountered inputs as familiar without recalling detailed stimuli information, which supports adaptive behavior across various timescales. We present a spiking neural network model with lateral connectivity shaped by unsupervised spike-timing-dependent plasticity (STDP) that encodes familiarity via local plasticity events. We show that familiarity can be decoded from network activity using both frequency (spike count) and temporal (spike synchrony) characteristics of spike trains. Temporal coding demonstrates enhanced performance under sparse input conditions, consistent with the principles of sparse coding observed in the brain. We also show how connectivity structure supports each decoding strategy, revealing different plasticity regimes. Our approach outperforms LSTM in temporal generalizability on the continual familiarity detection task, with input stimuli being naturally encoded in the recurrent connectivity without a separate training stage.

## Introduction

The brain can efficiently encode and later retrieve information about the stimuli encountered at various distances in time. Two processes contribute to the recognition memory: recollection and familiarity (Yonelinas, 2002). Recollection is oriented at the retrieval of information about the stimulus from memory, whereas familiarity is a simpler process of detecting whether a stimulus has been encountered before, via matching to existing representations. Familiarity memory operates in a temporally-agnostic fashion: familiar stimuli can be recognized at various distances in time within a certain timescale. On different timescales, familiarity can play a role in various processes: remembering things from childhood in lifelong learning, recognizing previously learned categories in continual learning tasks, and solving the exploration-exploitation dilemma within a single task. Low stimulus familiarity can facilitate its encoding as a new memory, or vice versa, its recollection if the stimulus is familiar (among others, Yonelinas et al., 1999; Yonelinas, 2001; Bülthoff & Newell, 2006).

Neural correlates of stimulus familiarity have been observed at different levels of visual hierarchy in animal studies. They range from stimulus-selective response potentiation to familiar stimuli in V1 (Cooke et al., 2015; Hayden et al., 2023), to repetition suppression in V2, inferior temporal cortex and perirhinal cortex (Brown et al., 1987; Miller et al., 1991; Ringo, 1996; Xiang and Brown, 1998; Anderson et al., 2008; Meyer & Rust, 2018), and even prefrontal cortex (Rainer and Miller, 2000). Computational models for familiarity recognition are based on a feedforward architecture with Hebbian or Anti-Hebbian learning (Androulidakis et al., 2008; Bogacz & Brown, 2003; Normal and O’Reilly, 2003). In Tyulmankov et al. (2023), a feedforward ANN with Hebbian plasticity was first shown to outperform LSTM in a continual familiarity task. Li and Bogacz have recently implemented recognition memory with use of an energy-based approach in Hopfield Networks and Predictive Coding Networks (2023).

However, limited attention has been given to time-agnostic familiarity encoding and detection in spiking neural networks. We show how familiarity can be encoded in lateral (recurrent) connections of a spiking network through fast unsupervised plasticity (Lazar et al., 2009). The decoding of familiarity is performed directly from the firing activity, namely from its both frequency (spike count in response to a stimulus) and temporal characteristics (spike synchrony, as first shown in in Korndörfer et al., 2017; Zemliak et al., 2024). We demonstrate that such a spiking network outperforms the trained LSTM in terms of time invariance, i.e. recognizing stimuli that were encountered at various intervals in the past. Additionally, we show that frequency- and synchrony-based familiarity decoding strategies require differences in the plasticity mechanisms.

## Results

### Encoding familiarity in lateral connectivity

The main mechanism for continuously encoding incoming stimuli in lateral connections of our network is a symmetrical STDP, which is also commonly referred to as Hebbian rule − unsupervised coincidence-based learning of local input features (Eq. 1-2). The algorithm utilizes the activity trace parameter, which allows for updating the connections for both neurons which fired recently, as well as a longer time ago.

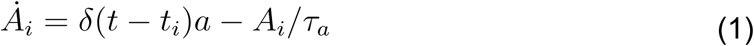

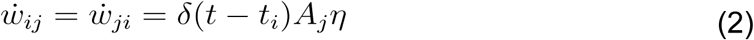

Eq. 1 describes the dynamics of activity trace *A_i_* of neuron *i*. *δ* is a dirac delta function, which defines spiking behavior of neuron i at the time moment t. *δ* = 1 if the neuron fires, and 0 if it does not. *τ_a_*= 20 ms is the trace memory parameter, which regulates how long the information about the activity of individual neurons remains in memory and can contribute to connectivity updates. *a* stands for an individual activity trace increase: its greater values mean stronger effect of firing events on weights.

Following Eq. 2, the plasticity update of a neuron’s lateral connections after firing is determined by the trace *A_j_* of every connected neuron. The resulting value is multiplied by the STDP update scaling factor *η*, which essentially is the learning rate of a model. Note that STDP updates between neurons in our model are symmetrical (Fig. 1C), due to the fact that the input stimuli are encoded in firing rates, rather than spike order, hence it is more important to emphasize the neurons which encode a single stimulus, and not their temporal firing pattern. We further discuss the input form in detail (see Continual familiarity experiments).

**Figure 1.**
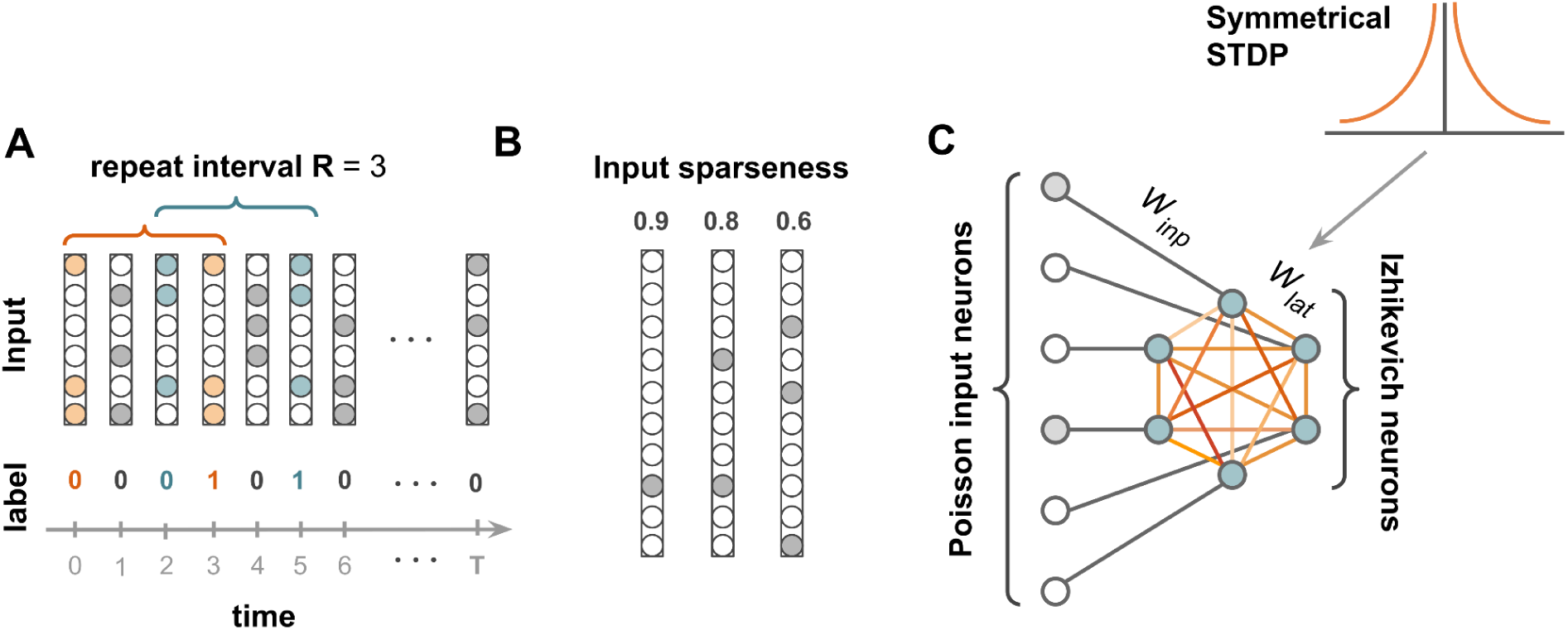
The dataset and the model. **A**. The structure of the dataset: binary vectors are organized along a temporal scale. Some of the vectors repeat after R time steps, others are randomly generated. Repeated vectors are considered familiar and are labeled as 1, non-repeated vectors are non-familiar and labeled as 0. **B**. Three degrees of input sparseness are used in the experiment: 0.6, 0.8, 0.9. **C**. The model architecture: each Izhikevich neuron has a one-to-one connection to the spiking input. Izhikevich neurons are laterally connected to one another. Lateral connections undergo symmetrical STDP.

Symmetric STDP can create a positive feedback loop in a laterally-connected network, whose weights are constantly growing and reinforcing neurons to increase their firing rate, which eventually leads to an excessive activity of a network. In our model, we use the synaptic weight normalization (Eq. 3) as a homeostatic mechanism which prevents the endless growth of connections and runaway dynamics (Gerstner & Kistler, 2002, Toutounji & Pipa, 2014). Weight regulatory mechanisms are of particular importance in a fully-excitatory network, since it does not have inhibition as an alternative way of stabilizing network activity.

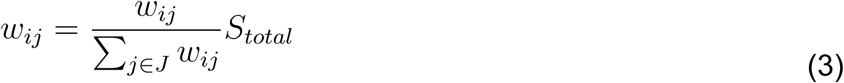

Here, *w_ij_* stands for a connection between neurons *i* and *j*. *J* is a subset of neurons which form incoming lateral connections to a neuron i. S_total_ defines the sum of all incoming lateral connections of a single neuron. It controls the overall activity level in the network.

In the context of input representations within the lateral connectivity matrix, the homeostatic mechanism is aimed at scaling the continuous connectivity updates and embedding them into the internal representation space. It has the additional parameter: normalization interval, which regulates how often normalization events happen, and thus how quickly new memories are rewriting the old ones. Longer normalization interval allows for more plasticity updates to increase the weights before they become rescaled, reducing existing old memories.

### Continual familiarity experiments

The network performance was evaluated on the continual familiarity dataset adapted from (Tyulmankov et al., 2022). The dataset is a sequence of 500 stimuli. Every stimulus is a 100-dimensional binary vector. The dataset is generated as follows: each stimulus is either a copy of the stimulus present in the dataset R steps ago, or a new randomly generated stimulus (Fig. 1). We refer to R as a repeat interval. R is the amount of new stimuli between two familiar stimuli in the dataset. The stimulus can be new with a probability p, or familiar, i.e. a copy, with a probability 1-p. For our experiments, p was set to 0.5. Additionally, each stimulus is characterized by its sparseness, i.e. the fraction of 0s in a binary 100-dimensional vector. In our experiments, we used stimuli with sparseness 0.6, 0.8, and 0.9 (see Methods Continual familiarity dataset). Intuitively, sparseness is the opposite of vector denseness: fewer 1s make a vector less dense and hence more sparse.

Every element of the stimulus binary vector represents an external input to a single neuron in the model (see Fig. 1). The binary stimuli vectors are transformed into a spiking input via the Poisson point process with the firing rate 100Hz for 1s in the vector, and 0Hz (no spiking) for 0s in the vector. Thus, whenever the stimulus is encountered for the second time in the dataset, i.e. is familiar, it is again generated from the same binary vector via the Poisson process. Thus, familiar stimuli are represented by entirely different temporal patterns: only their rate component is preserved.

The task is as follows: a network is presented with one stimulus at a time, and it has to predict whether the stimulus is novel or familiar. Our model operates continuously over time, with constantly ongoing unsupervised plasticity. We simulate 1000 ms of firing activity, and predict the stimulus familiarity from resulting spike trains. Note that familiarity can be estimated on longer intervals as well, and the accuracy either stays the same or even increases. We empirically selected the detection timespan which provided a tradeoff between the accurate and fast classification. The performance in all experiments was measured as prediction accuracy balanced by a proportion of novel stimuli in the dataset (see Methods Measuring model performance).

We used a genetic algorithm to preliminarily optimize STDP (activity trace memory, activity trace increase, STDP update scaling factor) and homeostatic plasticity (total incoming lateral weight, weight normalization interval) meta-parameters used in the experiments (see Methods Meta-parameter optimization). The parameters were optimized for different R values (3, 5, 10) and input sparseness levels (0.6, 0.8, 0.9; see Methods Continual familiarity dataset), separately for frequency- and synchrony based familiarity decoding methods. After optimization, we conducted a series of simulation experiments, to test model capabilities for continual familiarity detection on R values from 1 to 30. This allowed us to evaluate model generalizability over time: whether it can detect familiar stimuli on various time distances in the past, even if optimized for a specific distance only. This generalizability was evaluated separately for spike synchrony and spike count decoding methods, and for different input sparseness levels.

For our experiments, we used LESH – a laterally-connected excitatory spiking neural network with Izhikevich model dynamics (Izhikevich, 2003) and Hebbian-type STDP learning mechanism. Our network consists of a single hidden layer of 100 spiking neurons with all-to-all lateral connectivity (Fig. 1). Every neuron in the hidden layer follows Izhikevich dynamics, which describes how its membrane potential evolves over time (see Methods Izhikevich spiking model). When the membrane potential exceeds a firing threshold, a spike event is registered. The membrane potential is influenced by incoming spikes, both from the external input and lateral connections from other Izhikevich neurons.

### Continual familiarity classification

In our study, we used two methods for classifying, or decoding stimulus familiarity from the network activity: spike count and spike synchrony. Both are simple threshold methods: we measure a certain parameter of the model output spike traces, and find a threshold for the prediction. The threshold for classification is determined through preliminary genetic optimization (see Methods Meta-parameter optimization). Stimuli with the measurement value below the threshold are identified as novel, and stimuli with above-threshold values – as familiar.

The first decoding method is a spike count. This is a frequency method which does not depend on the temporal structure of output spike trains. Another metric is called Rsync (Eq. 11), and it estimates spike synchrony in a fully-excitatory network, following (Korndoerfer et al., 2017; Zemliak et al., 2024).

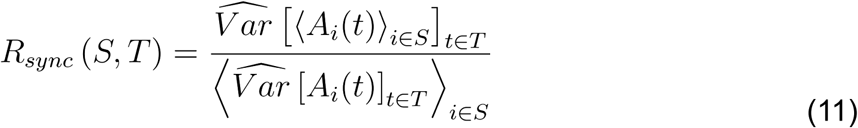

Here, S defines the neuron population, and *T* is a time interval for measuring synchrony within a given interval. It is calculated as the variance of the average activation trace *A_i_* of the neural population *S*, divided by the average of individual variances of neurons from the population. *A_i_* represents an activation trace of the neuron *i* from population *S*. It is calculated as follows: a raw binary spike train of the neuron *i* is convolved with an exponential kernel *k(t)* = *e^-2t^*, with the timescale = 3 ms. In related studies estimating spike synchrony, the timescale varies from 1 to 10 ms (Pipa et al., 2008). We chose a timescale from this range which is comparable to the timescale of an EPSP (Pipa et al., 2008). As shown in (Zemliak et al., 2024), the precise choice of a timescale does not have an impact on the result of Rsync measurement.

For both measurements, only spike trains from the neurons which fired at least once were used. The reason is, Rsync can be sensitive to the amount of spike trains, and we decided to fix the amount of neurons in the measure. We measured average spike count in a similar fashion, to be able to compare the performance when optimized for every decoding measure.

### LESH generalizes across repeat intervals

To evaluate the performance of the LESH model, we compared it to the performance of LSTM (Hochreiter and Schmidhuber, 1997) and HebFF (the infinite-data experiment from Tyulmankov et al., 2022). LSTM is typically used in machine learning for solving memory tasks. HebbFF is utilizing a similar mechanism for memorizing stimuli as LESH - fast Hebbian plasticity. It was also recently used for solving a continual familiarity task, and overperformed LSTM (Tyulmankov et al., 2022).

We trained LESH, HEbbFF and LSTM for individual R values 3, 6 and 10, and then evaluated their performance for the other R values from 1 to 30 (Fig. 2). Both LESH and HebbFF plastic weights were continuously updated in the experiments, while LSTM weights were fixed after training. LESH and LSTM models were evaluated on data of sparseness 0.8, and HebbFF on the data of sparseness 0.0. HebbFF encodes inputs as binary vectors of −1 and 1, which is not applicable to different sparseness values. LESH and HebbFF demonstrated better generalizability than LSTM, although LSTM performed superiorly on R used for optimization.

**Figure 2.**
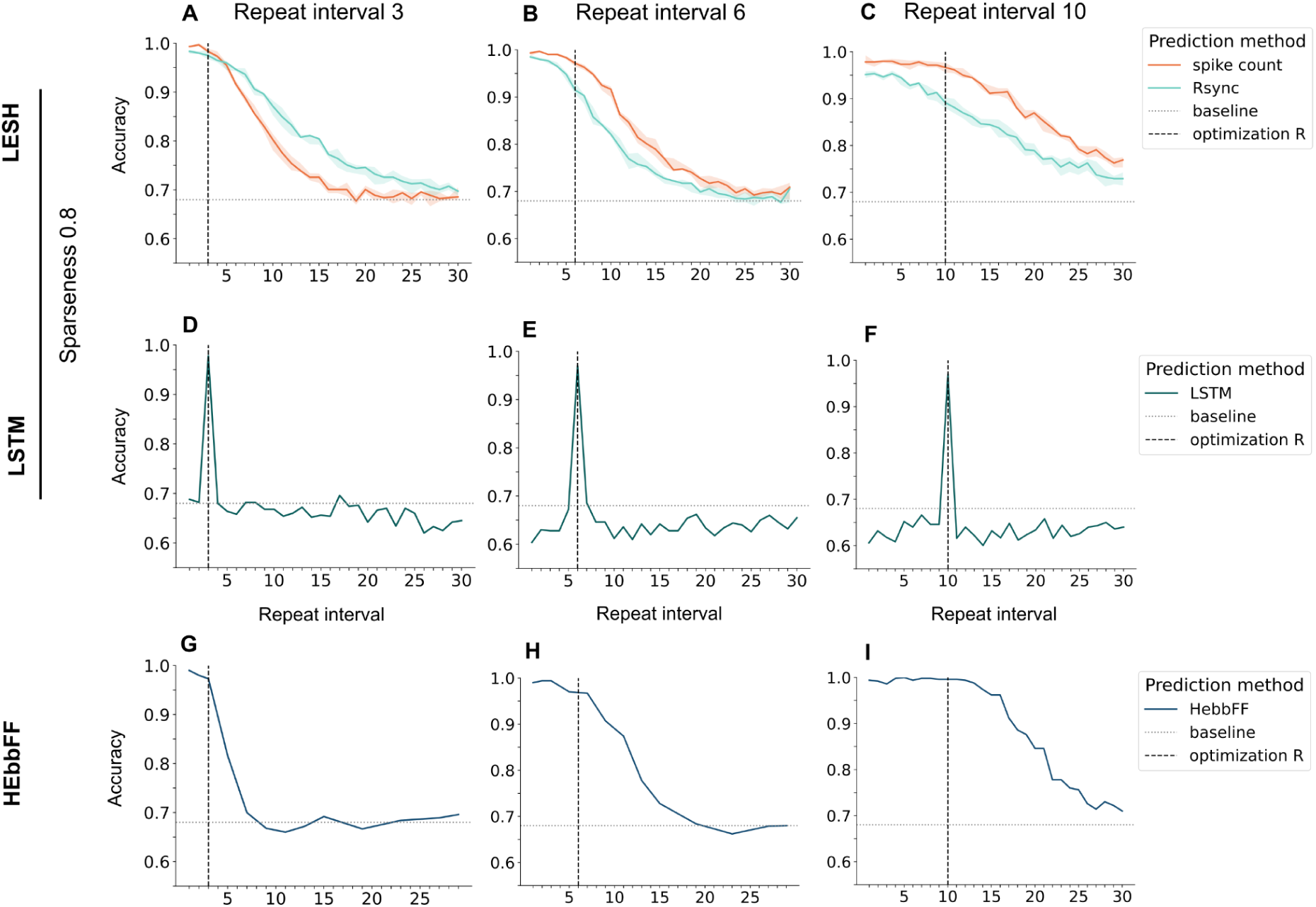
Models generalization across repeat intervals on the continual familiarity task. LESH, HebbFF, and LSTM models evaluated on datasets with different R values after a single optimization procedure each. LESH and LSTM optimized for sparseness 0.8, HebbFF for non-sparse stimuli. A baseline corresponds to always selecting the most frequent class. **A-C**. The LESH model with STDP parameters optimized for a specific repeat interval R, is evaluated on different R values on the continual familiarity task. LESH always extrapolates over the repeat interval that its learning parameters were optimized for. **D-F**. The LSTM model trained on a specific R value does not extrapolate at all, when evaluated on the other R values. **G-I**. The HebbFF model optimized for a specific R in the infinite-data regime (see Tyulmankov et al., 2022), can extrapolate over the other R values.

For the LESH model, both spike count and spike synchrony classification strategies led to a high generalizability across R values, but spike synchrony had higher accuracy. Information about stimulus familiarity is encoded in both frequency and temporal characteristics of the spikes, but the spike count measure demonstrates higher accuracy in all of the conditions and thus is more practically applicable for input of the sparseness level 0.8. STDP allows LESH to naturally remember stimuli encountered more recently than a target R steps ago, and even further in the past than R steps. For both spike count and synchrony, LESH performance gradually decreases with the larger R values, although the overall performance is noticeable higher for the spike count. Thus, although the model learning parameters were preliminarily optimized for a specific repeat interval via the genetic algorithm (see Methods Meta-parameter optimization), LESH can flexibly access memories on various distances in time.

When compared with HebbFF, the latter demonstrates either similar (for R values 3 and 6) or better (for R equal 10) performance for the repeat interval it was optimized on, however its performance on the other R values decreases more rapidly than the performance of LESH. This is especially illustrative on the repeat interval 10: HebbFF demonstrates slightly better performance for R below 15, but then has a steep decline in its generalization abilities. The LESH performance, however, declines smoothly with an increase of R, which shows its good generalization capability.

We also compared the performance of LESH and LSTM across input sparseness values.The performance of LSTM did not depend on sparseness, so we only report the results for LESH. It performed better for sparser stimuli, but demonstrated the ability to generalize over R, for all sparseness values (Fig. 3).

**Figure 3.**
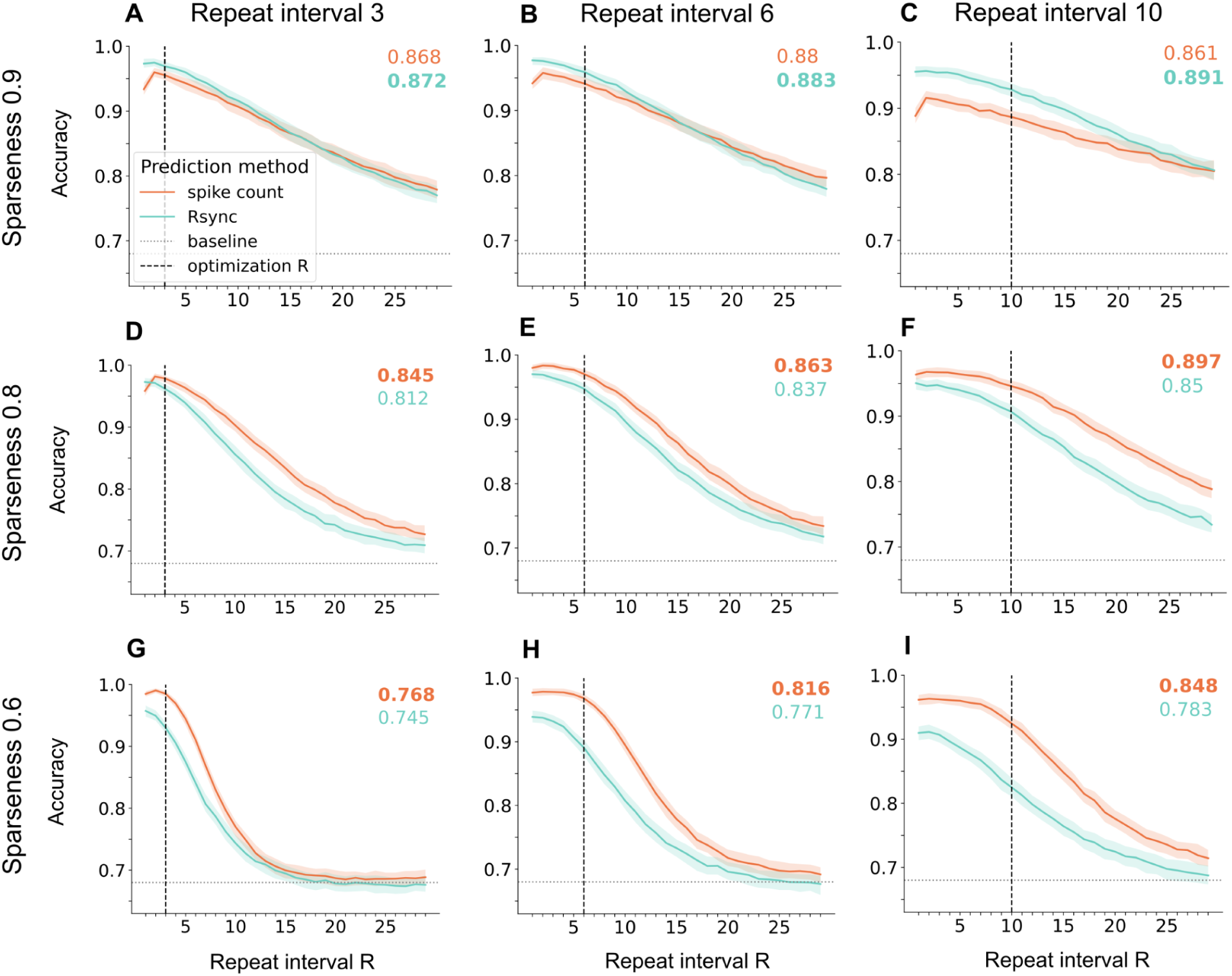
LESH performance on data of various sparseness. The model performs better on sparser input stimuli. Colored numbers reveal generalizability: accuracy averaged across all test repeat intervals (a larger number in **bold**). To ensure stability of the results, models were independently optimized for 20 times and yielded 20 parameter sets for every combination of sparseness and repeat interval values. The resulting plots were received after averaging accuracy for each combination. **A-C**. LESH performance for data with sparseness 0.9. **D-F**. LESH performance for data with sparseness 0.8. **G-I**. LESH performance for data with sparseness 0.6.

LESH ability to generalize over repeat intervals highly depends on sparseness of the data, as well as on the repeat interval that its plasticity parameters were optimized for. The overall performance of our model indeed drops with decreasing sparseness of the input stimuli. The reason is, a certain amount of internal representations needs to be stored in a network simultaneously, depending on the input sparseness and the repeat interval. These representations can overlap and make the familiarity detection more difficult. Consider a novel stimulus which overlaps a lot with internal representations of familiar stimuli already encoded in a lateral connectivity matrix. The overall activity in response to such a stimulus can be indistinguishable from the response to a familiar stimulus.

### Plasticity regimes regulate representations learning

The model performance depends on its ability to distinguish new memories from the representations already encoded in lateral connectivity, and hence on the ability of plasticity to encode these representations in a reliable manner for subsequent readout. Thus, STDP meta-parameters shape the connectivity structure, which in turn guides the interplay between neurons in the network through lateral connections. The optimal parameter combination allows for the connectivity, which induces maximally different responses of a network to novel and familiar stimuli.

All plasticity meta-parameters in LESH were optimized using a genetic algorithm (see Methods Meta-parameter optimization) for a specific combination of repeat interval and input sparseness, thus they reflect how STDP adapts to different memorization needs. The plasticity has to learn to distinguish between overlapping memory representations for stimuli of low sparseness, and to keep more representation in memory simultaneously for longer repeat intervals. To analyze how plasticity shapes connections in a network, we computed Pearson correlations between its meta-parameter values and several characteristic features of the connectivity matrix, such as Gini index, modularity, transitivity, participation coefficient and betweenness-centrality (Fig. 4). These metrics describe clusteredness of the network, i.e. sizes, shapes and overlap of memory representations.

**Figure 4.**
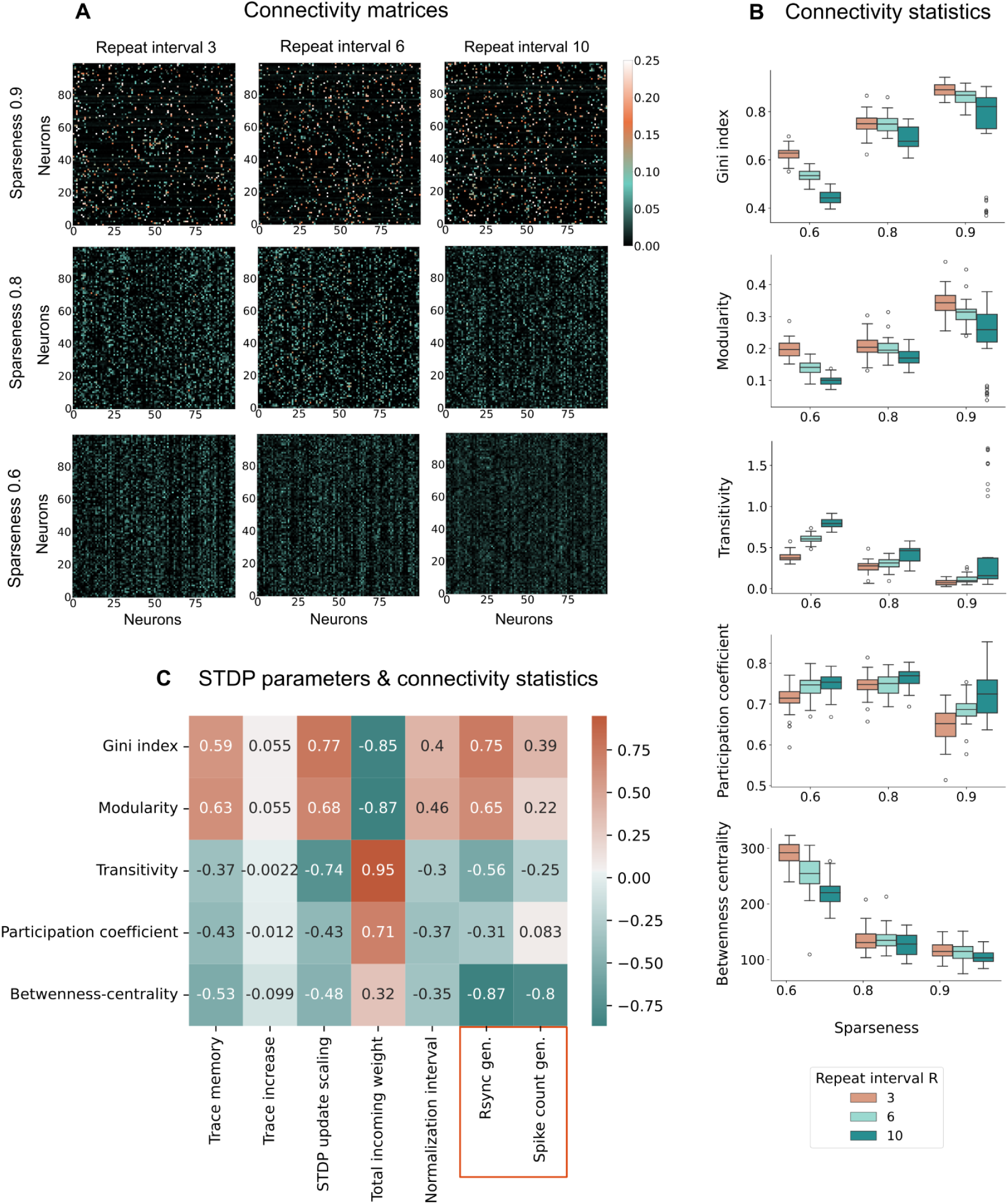
Connectivity adaptation. Various connectivity statistics of the optimized models. Data is presented for the LESH model optimized for spike count, after continuously encountering 500 input stimuli of various sparseness 20 times, for 20 independently optimized parameter sets, each corresponding to a repeat interval+input sparseness values combination. Note that statistics remain similar for the parameters optimized for Rsync. Weights were preliminarily rescaled via division by the lateral connection sum values, optimized for each sparseness level and repeat intervals (see Eq. 3). **A.** Connectivity matrices structure across input sparseness levels and repeat interval values. **B**. Correlation of STDP parameters with connectivity measured features, presented for sparseness level 0.8 and repeat interval 6. Sparser inputs result in sparser connectivity, expressed in higher modularity, higher Gini index, lower transitivity, participation coefficient and betweenness-centrality. For highest (0.9) and lowest (0.6) sparseness values, connectivity statistics also differ across repeat intervals. **C**. Pearson pairwise correlation between connectivity structure statistics and plasticity meta-parameter values + Rsync and spike count generalizability values.

In the context of our network, Gini index and modularity describe how well it can be divided into distinct individual representations encoded in lateral connectivity, i.e. how strongly separate representations overlap. Sparse stimuli lead to more heterogeneous connectivity, with groups of strong connections among many weak ones, which is reflected in higher Gini index and modularity (see Fig. 4A-B). Gini index and modularity also have the smallest values for long repeat intervals, which corresponds to more representations being encoded in the connectivity at the same time.

Gini index and modularity strongly correlate with trace memory and STDP update scaling factor (see Fig. 4C), which helps to sharpen the plasticity updates between the neurons encoding one representation, in relation to connections to neurons outside of it. These parameters have a stronger positive effect on the performance of Rsync than spike count in a task of familiarity detection, which suggests that Rsync is more sensitive to the representations independence from each other. This explains Rsync better performance on more sparse input stimuli, which lead to a formation of more distinct representations.

In comparison, the total incoming weight parameter substantially increases the magnitude of connections between neurons belonging to different representations, which negatively impacts the modularity of the connectivity structure. This is an inevitable consequence of the all-or-none nature of neuron firings: STDP is based on the spike coincidence detection, and the weight of any individual neuron cannot be reduced further than a certain level, in order for it to be able to trigger other neurons to fire and hence form meaningful connections with them. Thus, the more neurons are encoding a single representation of the least sparse stimuli, the larger incoming weight parameter is required for STDP to encode representations necessary for further classification.

Participation coefficient estimates the representations overlap: it measures how much neurons are involved in multiple tightly clustered representations. Interestingly, it negatively impacts Rsync performance, but not spike count. This falls in line with the performance differences of Rsync and spike count across sparseness levels (see Fig. 3). Due to the emergent nature of Rsync, a synchronous volley of activity can distribute through the neurons with high participation coefficient onto the tightly connected neurons encoding a “false” representation. Unlike participation coefficient, high betweenness-centrality signals about dense between-cluster connectivity not through several highly connected overlapping neurons, but rather many weakly connected ones. This severely disrupts both Rsync and spike count familiarity detection.

### Spike synchrony and spike count require different plasticity regimes for inputs of various sparseness

The optimal STDP regimes not only vary greatly for different sparseness and repeat interval values, but also differ for familiarity classification based on spike synchrony and spike count (Fig. 5). The plasticity regime is defined by meta-parameters that control various aspects of the plasticity process, such as the size and the frequency of weight updates, the longevity of activity traces, etc. Different input conditions and classification strategies lead to best performance under different STDP parameters, which are found during the optimization.

**Figure 5.**
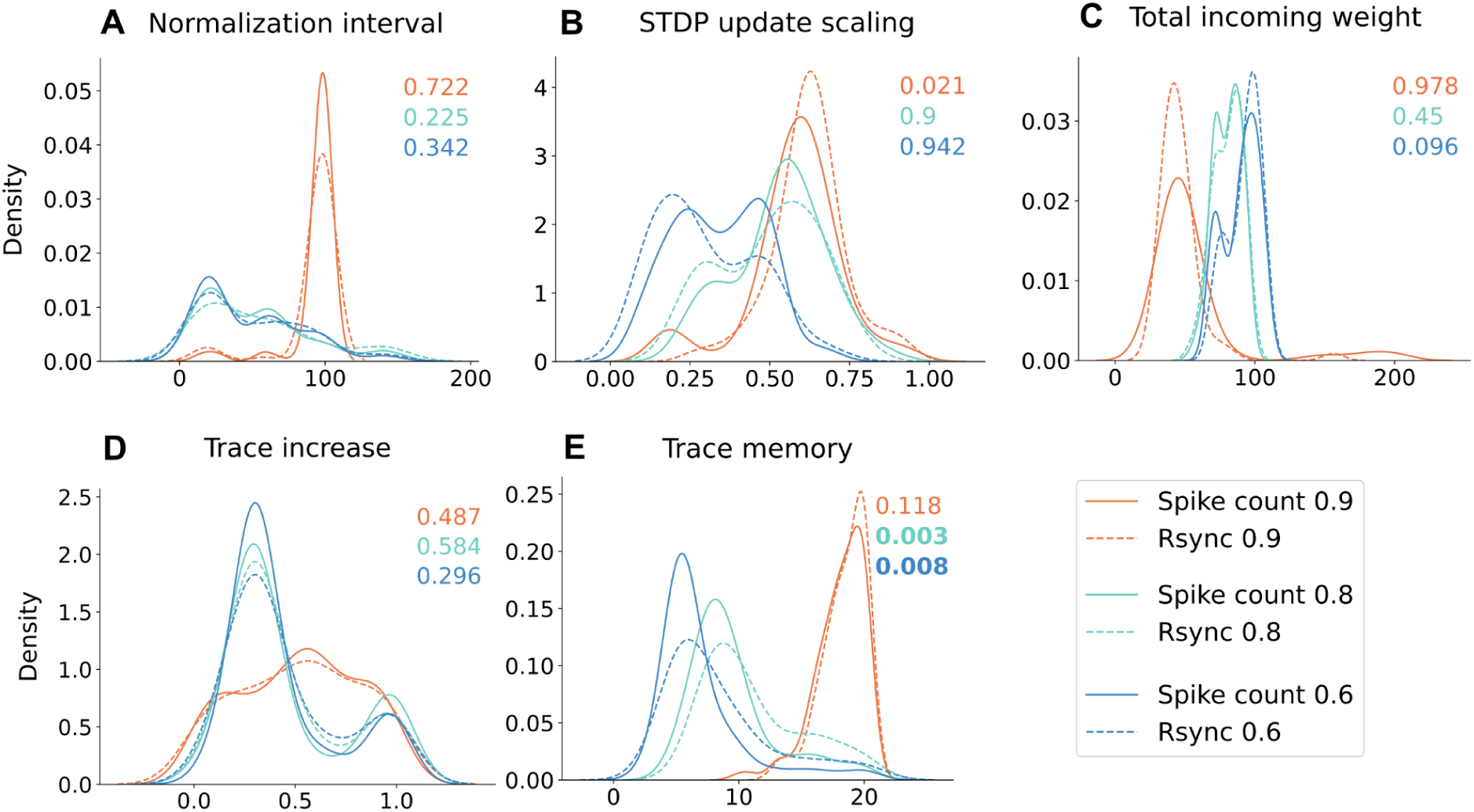
Optimal parameter distributions across sparseness levels. Parameter distributions more strongly differ across sparseness levels than familiarity detection measures (Rsync and spike count). Each distribution, represented by a colored curve, includes 20 parameter values received during 20 independent optimization procedures. Color stands for sparseness: orange 0.9, green 0.8, blue 0.6. Colored numbers stand for p-values for differences between measures within every sparseness level, computed via a permutation test with 10000 permutations and Bonferroni correction for multiple comparisons. Data for plasticity meta-parameters: **A**. Normalization interval, **B**. STDP update scaling factor, **C**. Total incoming weight, **D**. Trace increase, **E**. Trace memory. Differences across measures are significant (p < 0.05) for E within the sparseness levels 0.6 and 0.8.

In general, all parameters which account for plasticity weight updates, namely STDP update scaling factor, trace increase, and trace memory, are higher for the most sparse and thus last overlapping stimuli. Greater updates naturally lead to better memorization and thus easier further familiarity detection: both spike count and spike synchrony threshold measures work most efficiently, when novel and familiar stimuli yield maximally different responses. We purposefully fixed the value of external input scaling factor (see Table S1), thus the activity level in response to novel stimuli cannot be regulated much. During the optimization procedure, LESH had to identify the parameter sets which allow for maximal possible increased activity in response to familiar stimuli.

However, less sparse stimuli impose constraints on STDP weight update parameters, for two reasons. First, since less sparse input means more neurons receiving the external input encoding a single stimulus, the overall activity in the network has to be regulated. Second and most interestingly, when new and existing representations start to overlap for less sparse stimuli (see Fig. 4B), novel and familiar stimuli might evoke falsely similar responses. Thus, weight updates have to be adjusted, which leads to decreased parameters regulating STDP updates, for less sparse stimuli. This markedly impacts spike synchrony, which requires stronger weights within a single representation to perform efficiently.

Trace memory is the only parameter that differs significantly between spike count and spike synchrony optimizations. It represents how long the activity traces of stimulus-encoding neurons remain strong *during the presentation of a single stimulus*. Longer trace memories allow neurons involved in a specific memory to form stronger connections with each other. This difference becomes significant for inputs with sparseness levels of 0.6 and 0.8, where the inputs overlap and, as a result, their internal representations in the network also overlap. Synchrony in a network depends on strong and even connectivity to quickly spread activation and make neurons fire coherently in time. When the overlap between internal representations increases at sparseness levels of 0.6 and 0.8, the Rsync optimization uses longer trace memory to enhance synchrony within each familiar memory.

The total incoming weight is greater for less sparse stimuli because the number of neurons encoding each representation doubles at each sparseness level (see Methods: Continual Familiarity Dataset). For STDP, sufficient lateral input is needed to trigger firing, as memory representations form when neurons fire together and reinforce each other. Thus, the total input across all neurons in a representation increases.

The repeat interval used for optimizing plasticity parameters affects model performance and connectivity, though less significantly (Fig. 6). Longer repeat intervals require parameters that can reliably encode more stimulus representations for future recall. Input sparseness determines how many neurons store each stimulus, while the repeat interval defines how many representations must be stored simultaneously. The STDP update scaling factor is the only parameter with notable differences between repeat intervals: its value is lower for the longest repeat interval 10. This slows weight updates, allowing older representations to be rewritten more gradually and remain available for longer. This enables the network to recognize stimuli encoded over longer intervals, improving generalizability.

**Figure 6.**
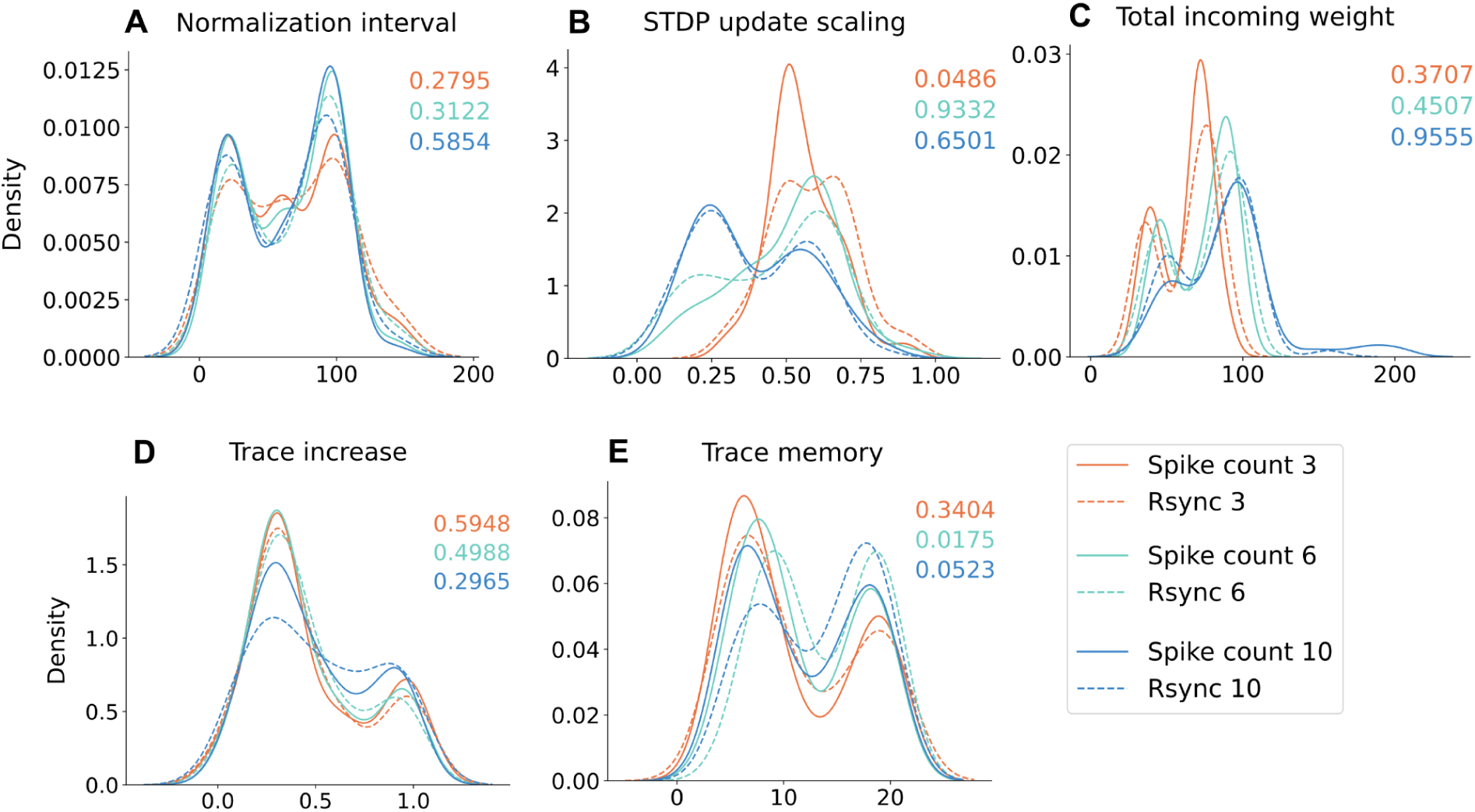
Optimal parameter distributions across repeat intervals. Parameter distributions more strongly differ across optimization R than familiarity detection measures (Rsync and spike count). Each distribution, represented by a colored curve, includes 20 parameter values received during 20 independent optimization procedures. Color stands for R: orange 3, green 6, blue 10. Colored numbers stand for p-values for differences between measures within every R, computed via a permutation test with 10000 permutations and Bonferroni correction for multiple comparisons. Data for plasticity meta-parameters: **A**. Normalization interval, **B**. STDP update scaling factor, **C**. Total incoming weight, **D**. Trace increase, **E**. Trace memory. No significant differences between parameters for different familiarity measures.

Along with the individual parameter values, we also investigated the relation of the model’s plasticity parameters to each other (Fig. 7). The interplay between the parameters is also important for adjusting the plasticity mechanism between synchrony and spike count-based regimes of familiarity detection. A synchrony-focused regime, i.e. the regime where the familiarity detection is performed based on the level of synchrony, requires stronger memorization in the form of longer memory traces, and thus requires more balancing for the enhanced activity during memorization.

**Figure 7.**
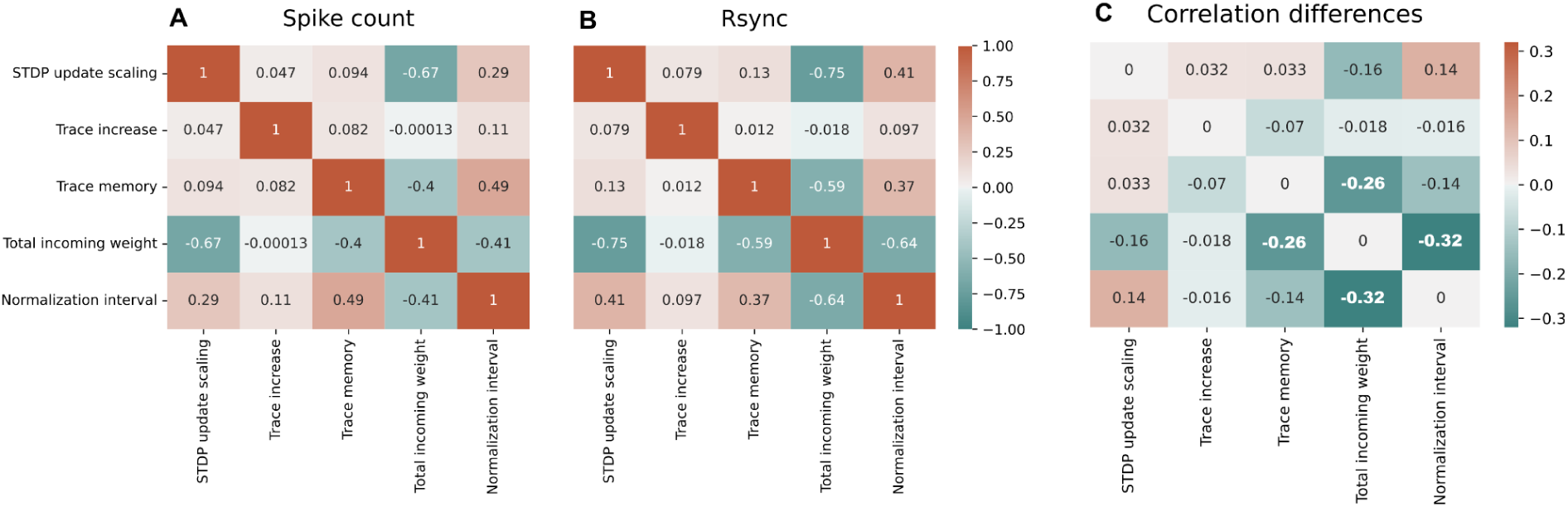
Correlations in parameter sets optimized for Rsync and spike count. Pairwise correlations for parameters optimized for: **A**. Rsync, **B**. Spike count. **C**. Differences between Z-transformed Pearson correlation scores from A and B. Every correlation score is averaged across 20 parameter value pairs received during 20 independent optimization procedures. **Bold** font stands for p < 0.05 (total incoming weight & trace memory, normalization interval). P-values were calculated using a permutation test with 1000 permutations, with Bonferroni correction for multiple comparisons.

Although the correlation matrices are relatively similar for spike count and spike synchrony, all negative correlations are stronger for spike synchrony. Negative correlations across all parameter sets exist only between total incoming weight and all other plasticity parameters, since it increases between-representation connections to a level where they start interfering with the modular structure of the network (see Plasticity meta-parameters shape representations learning). Moreover, only differences in two negative correlations of the total incoming weight, namely with trace memory and normalization interval, are of significance. As discussed above, synchrony requires more solid memorization of a stimulus than spike count to perform efficiently, which is realized via longer memory traces and intervals between normalization events. Such enhanced synaptic plasticity is balanced through the weight scaling (a smaller total incoming weight), to prevent excessive activity in the network.

In sum, these correlations illustrate the trade-offs in optimal fast plasticity regimes for LESH networks across different input encodings, optimization goals, and preferred read-outs. Sparse encodings with well separated representations lead to the best performing LESH networks but require stronger lateral connectivity to sustain sufficiently strong activity in the network to support effective fast plasticity. Optimizing for longer repeat intervals can also increase generalised performance (Fig. 3) and requires fast plasticity to have a stronger effect on connectivity to store more patterns. Similarly, optimizing for a synchrony read-out instead of rate requires longer activity traces to strengthen connections between co-activated neurons. Weight normalization emphasizes this effect.

### Predicting LESH performance from the connectivity features

Different plasticity regimes and input encodings lead to distinct patterns of network connectivity. This raises the question of whether the specific features of this connectivity predict how effectively the network responses distinguish familiar from unfamiliar stimuli. Therefore, we analyzed various connectivity features and their combinations to identify those with the greatest influence on LESH performance in a familiarity detection task. To achieve this, we trained and evaluated simple Decision Tree and Linear regression models using a 10-fold cross-validation procedure (see Methods Regression analysis for details). These models were then used to predict generalized performance of LESH in a continual familiarity task via spike count or synchrony, based on connectivity features (see Table 1).

**Table 1.** Predicting generalizability from connectivity.

| Regression method | R2 | RMSE |  | Gini index | Transitivity | Betweenness-centrality |
| --- | --- | --- | --- | --- | --- | --- |
| **Spike count** |  |  |  |  |  |  |
| Linear regression | 0.656 | 0.022 |  | 0.0 | 0.0 | -0.0004 |
| Decision tree | 0.67 | 0.019 |  | 0.136 | 0.092 | 0.772 |
| **Rsync** |  |  |  |  |  |  |
| Linear regression | 0.755 | 0.023 |  | 0.0 | 0.0 | -0.0005 |
| Decision tree | **0.845** | **0.017** |  | 0.149 | 0.033 | 0.818 |

Predicting model performance generalizability in a continual familiarity task, using Rsync and Spike count, from connectivity characteristics. The table includes R2 and root mean squared error (RMSE) regression performance metrics and feature importances/coefficients for Decision tree and Linear regression regression methods respectively (see Methods Regression analysis for details on feature importances calculation). Rsync performance can be predicted better from connectivity features, i.e. is more dependent on plasticity hyperparameters. Linear regression yields worse results than a non-linear Decision tree regressor, because predictor variables have complex non-linear interactions. Results in the table were received from 10-fold cross-validation, on 360 observations in total. See Methods Regression analysis for all hyperparameters and details on the regression procedure.

We used three characteristics of connectivity matrices for the regression analysis: Gini index, transitivity and betweenness-centrality, since all features were highly correlated with each other, and the addition of more than three features into the model had no impact on the performance. The particular set of predictor features was selected using grid search (see Methods Regression analysis). Both resulting performance metrics were substantially higher for the Decision Tree regression, suggesting that features have nonlinear interactions which cannot be captured by a linear regression model. Both models were able to better predict the Rsync than spike count generalizability value from connectivity features.

Betweenness-centrality was the most important for prediction feature, for both spike count and Rsync performances on a continual familiarity task. The predicted performance was negatively correlated with betweenness-centrality, which measures the representations overlap in a connectivity matrix. The Gini index played a bigger role in predicting Rsync performance compared to spike count. This falls in line with our observation that trace memory, which is reflected in the Gini index and modularity of the connectivity matrix, is the only plasticity meta-parameter which differs significantly between networks optimised for spike count and spike synchrony read-outs.

The result that performance of the Rsync network is very well predicted by network connectivity, and the result that prediction of the performance of the rate network is possible but significantly worse in comparison, highlights that spike synchrony is much more sensitive to features of network connectivity. This is in line with previous research showing how specific connectivity patterns lead to specific rsync responses (Zemliak et al., 2024) and may also help explain that robust performance was easier to achieve in rate networks.

## Discussion

We demonstrate that a laterally-connected spiking neural network with local Hebbian plasticity can successfully perform a continual familiarity task, and generalize across repeat intervals. Our result is consistent with previous research on continual familiarity in networks with Hebbian-like learning (Tyulmankov et al. 2022), although the said work is comparing Hebbian and anti-Hebbian plasticity. Here, we focus on Hebbian plasticity for the clarity of an argument. We show that a simple form of unsupervised Hebbian learning in a spiking network is naturally encoding input familiarity in lateral connectivity, without the explicit training and any top-down error or reward signal. This familiarity information can be read-out from the connectivity via spike count or spike synchrony.

The performance of our model on a continual familiarity task depends on three main factors: the sparseness of the input and the repeat interval used for the STDP parameters optimization, as well as a readout method for familiarity detection. The influence of these factors is reflected in the differences in the optimal parameter values. The input sparseness is mediated by multiple parameters regulating the size of STDP updates, which are in turn balanced by the weight normalization across the neurons. The adjustment to longer repeat intervals is realized through changes in the “severity” of increasing connections for new memories, simultaneously rewriting the old ones. Finally, the readout method shapes the connectivity organization: spike synchrony favors more homogeneous internal representations.

We investigate two mechanisms for decoding stimulus familiarity from the model spiking activity: spike count and spike synchrony. Spike count is a simple frequency metric, whereas spike synchrony relies on temporal characteristics of the spike trains. Both metrics are viable for detecting a stimulus familiarity, although spike count proves to be more precise and robust across all experimental conditions. However, synchrony outperforms spike count as a threshold classification measure for the most sparse inputs, which allow for almost non-overlapping representations. When the overlap is large, spike synchrony falls behind the spike count due to the inherent dependance on the strong and homogeneous representations. This property of synchrony falls in line with previous spike synchrony simulation studies (Korndörfer et al. 2017; Zemliak et al. 2024). The representations inevitably weaken to counteract the overlap, which disrupts synchrony. Spike count performance also decreases for the last sparse stimuli, but not as dramatically.

However, we do not see the model dependance on the input sparseness as a weakness, but rather as an inherent feature of a spiking dynamical system, which falls in line with biophysical observations of the brain. Neurons of sensory and motor cortices demonstrate highly selective and sparse responses to the input stimuli (Olshausen & Field, 2004). The activity of only a small amount of V1 neurons is required to reliably decode the perceived images (Yoshida & Ohki, 2020). Similarly sparse responses have been demonstrated by excitatory neurons of the primary auditory cortex (Liang et al. 2018). Thus, the model’s reliance on sparse input aligns with the principles of sparse sensory coding in the brain.

Neurons in our model are equipped with strictly defined receptive fields: each neuron receives an input from a single corresponding spiking neuron. This can be likened to a simplified model of V1 from (Zemliak et al. 2024), although we intentionally do not specify the angle orientation characteristics of the stimuli. Nevertheless, we also use the system with one-to-one input connections, and with lateral connections shaped by gradually acquired experience in the form of memorizing the incoming stimuli. Such architecture with local synaptic plasticity top does not reflect the complexity of real brain networks and does not replicate the familiarity signals from V2 (Huang et al., 2018), IT (Anderson et al., 2008; Meyer & Rust, 2018) or prefrontal cortex (Rainer and Miller, 2000). In the said brain areas, stimulus familiarity induces the reduced neuronal activity, the effect known as repetition suppression. According to (Tyulmankov et al., 2022), repetition suppression can be modeled with the anti-Hebbian rather than Hebbian plasticity. Our model equipped with Hebbian plasticity demonstrates the opposite behavior (repetition enhancement), which however can be likened to the neuronal response to familiarity in V1 (Cooke et al., 2015; Hayden et al., 2023): neurons in the primary visual cortex tend to increase their firing activity in response to previously experienced stimuli.

Last but not least, our results suggest that spiking networks can be useful in continual learning tasks, since the input familiarity can be directly read out from their activity without an explicit training procedure, in a continuous fashion. For example, consider a classical continual learning paradigm where a model first has to learn one task, and afterwards is trained on another one. The main challenge for the model is to learn the new task without forgetting the old one. In this case, spike synchrony, or spike count, or another more complex metric could be used for detecting whether a stimulus has been encountered before and thus belongs to the old familiar task, or it corresponds to a new task. It has already been shown that spiking networks can excel in continual learning problems, due to the natural sparseness of their activity and a range of easily implementable methods to push this sparseness even further (Antonov et al., 2022; Shen et al., 2024). Moreover, it was recently directly demonstrated that task familiarity estimation in spiking neural networks can facilitate the efficient reuse of existing representations for new tasks, leading to improved performance and decreased energy consumption (Han et al., 2024). Therefore, we believe that frequential and temporal characteristics of spike trains can naturally encode the input familiarity, and this is a crucial feature for various kinds of continual learning.

## Acknowledgments

The work was supported by funds of the research training group “Computational Cognition” (GRK2340) provided by the Deutsche Forschungsgemeinschaft (DFG, German Research Foundation). The high performance computing cluster used to run the experiments was also funded by the German Research Foundation (DFG) − 456666331.

## Author Contributions

Conceptualization, V.Z. and P.N.; Methodology, V.Z. and P.N.; Software, V.Z.; Validation, V.Z. and P.N.; Formal Analysis, V.Z.; Investigation, V.Z.; Resources, P.N. and G.P.; Data Curation, V.Z.; Writing – Original Draft, V.Z. and P.N.; Writing – Review & Editing, V.Z., P.N., and G.P.; Visualization, V.Z.; Supervision, P.N. and G.P.; Funding Acquisition, G.P.

## Resource availability

### Lead contact

All requests should be directed to and will be fulfilled by the lead contact, Viktoria Zemliak.

### Materials availability

This study did not generate new unique materials.

### Data and code availability

All data in the work was programmatically generated. The code for generating data, running simulations and analyzing the simulation results, as well as the simulated data logs, plots and statistics, can be found at

https://github.com/rainsummer613/spiking-continual-familiarity (https://doi.org/10.5281/zenodo.14639677). Any additional information regarding the data generation and analysis is available from the lead contact upon request.

## Methods

### Continual familiarity dataset

A network is continuously presented with a set of stimuli. Every time moment, one stimulus is presented, and this is considered to be a single dataset sample. A stimulus is a 100-dimensional binary vector consisting of 0s and 1s, and their proportion defines the sparseness parameter of the dataset. The more 0s and fewer 1s there are in a stimulus binary vector, the higher is the sparseness value (Eq. 5).

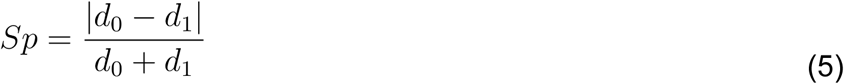

Here, *Sp* is a sparseness parameter of the stimulus. *d_0_*and *d_1_* define the amount of 0s and 1s in a binary stimulus vector respectively. Note that *d_0_ + d_1_* defines a dimensionality of the input vector, which also equals the amount of Izhikevich neurons in the model. In our experiments, we used three combinations of *d_0_* and *d_1_*values, corresponding to three levels of sparseness (se Table 2). A dataset consists of stimuli of the same sparseness.

**Table 2.** Stimulus sparseness.

| $d_0$ | $d_1$ | Sparseness ($Sp$) |
| --- | --- | --- |
| 80 | 20 | 0.6 |
| 90 | 10 | 0.8 |
| 95 | 5 | 0.9 |

The proportion of 0s and 1s in the binary stimulus vector defines sparseness of the stimulus in a dataset.

Each stimulus is randomly generated with probability 1-p, and with probability p it repeats the stimulus from the time step t-R, i.e. R time steps ago in the dataset. We restrict every stimulus to appear no more than 2 times within a single dataset, hence a single stimulus can only repeat once. We generate our dataset with a new stimulus probability 0.5, thus the fraction of familiar stimuli is 1/3, and the fraction of novel stimuli 2/3 respectively. The procedure for the dataset generation was first presented in (Tyulmankov et al., 2022).

In our setup, a single dataset has a fixed repeat interval R, and the model performance is evaluated on multiple datasets with R values from 1 to 30. Before the experimental simulations for performance evaluation, the model undergoes the optimization stage with a fixed R value (see Methods Optimization for details). In our experiments, R for optimization is fixed to either 3, 6, or 10. The evaluation is performed for multiple R values, to estimate the model generalization ability, as in (Tyulmankov et al., 2022).

### Hyperparameter optimization

In our spiking network, the Izhikevich neuron model parameters are fixed, and several Hebbian plasticity and connectivity parameters are subjects to the optimization, which takes place before the experimental simulations. The following parameters are optimized: lateral input scale, plasticity scale, total lateral input, weight normalization interval, and either spike count or Rsync classification threshold. In the present study, the optimization was performed with a variation of a genetic algorithm (Holland, 1992; Fig. 9).

**Figure 9.**
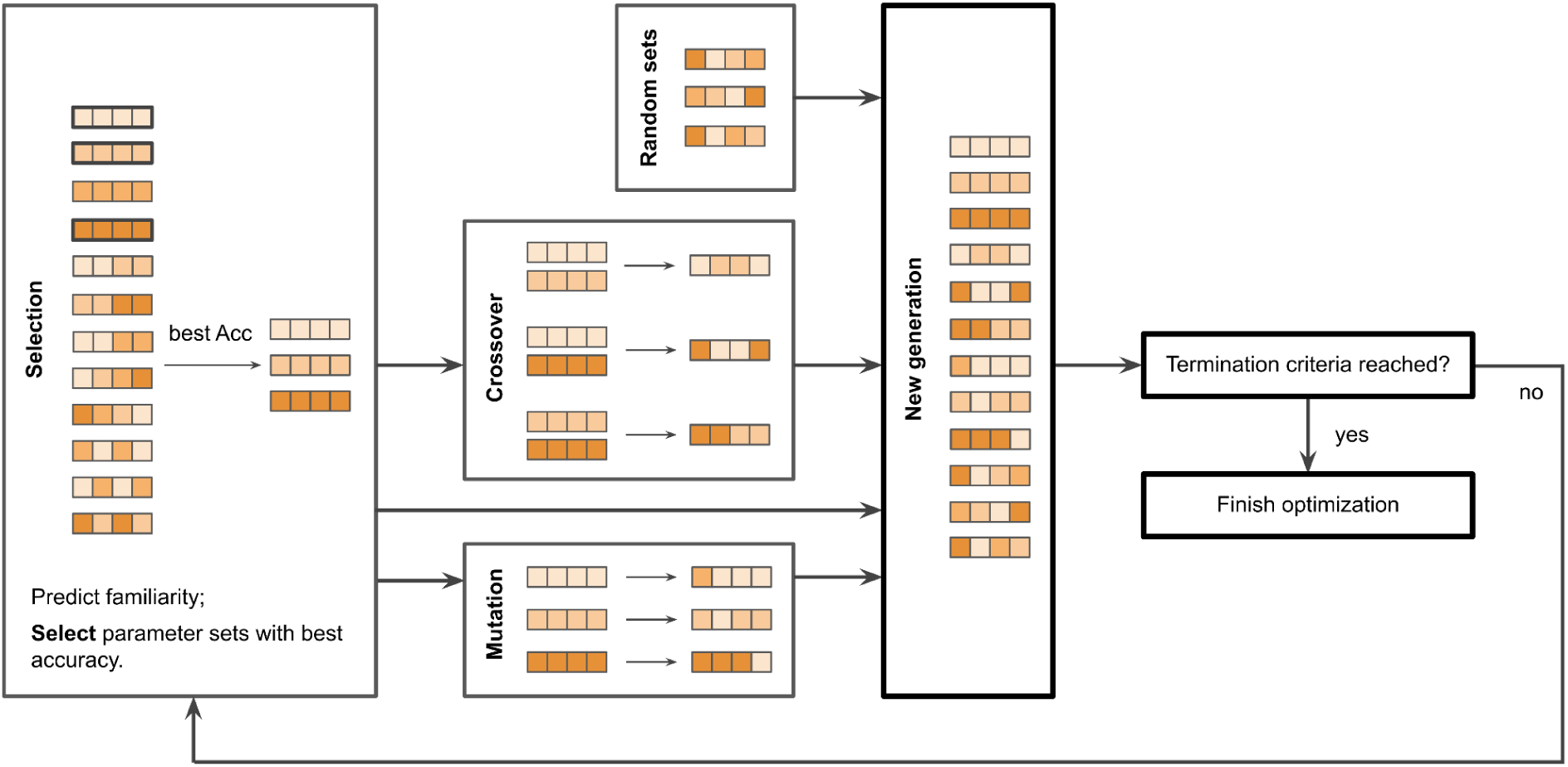
The genetic algorithm. A pipeline of the genetic algorithm. There are 12 parameter sets in every generation, i.e. in iteration of the algorithm. Spike trains are generated with an Izhikevich model and with each parameter set in the generation. Then, accuracy of familiarity detection is measured with use of spike count and Rsync decoders. Those parameter sets which lead to the best performance, are selected for the next generation of the genetic algorithm. They undergo operations of crossover and mutations; also, new parameter sets are generated. The algorithm finishes if any of the termination criteria are met: 200 generations are over; accuracy 1.0 is achieved for any parameter set in a generation; accuracy has not increased for 15 generations.

The algorithm was run for 300 generations, each generation has 12 descendants (i.e. parameter sets), and for every generation the following operations were performed:

1. Each parameter set is used to evaluate the model on a dataset of N=500.
2. The performance of the simulations with all parameter sets from the current generation is evaluated and compared. Note that we separately optimize the parameters for Rsync and spike count metrics.
3. Mutation: in each of 3 best-performing parameter sets, one random parameter is multiplied by a random factor from 0.75 to 1.25.
4. Crossover: from 3 best-performing parameter sets, 3 pairs are formed. In each pair, a random half of parameter values are taken from one set, and the rest – from another.
5. Generation: 3 new random parameter sets are generated.
6. Next generation of 12 parameter sets is formed from: 3 best-performing parameter sets of the current generation, 3 sets formed as a result of mutation, 3 sets formed as a result of crossover, and 3 randomly generated sets.

The algorithm described runs either 300 generations in a row, or until the accuracy of 1.0 is achieved by one of the parameter sets, or until in 15 generations in a row the best-performing set of the current generation performs worse or the same as the best-performing set of the previous generation. The classification threshold is determined at each generation iteratively, as the threshold which leads to most accurate classification. When the optimization is finished, the corresponding threshold is fixed for the subsequent experimental simulations.

Importantly, the optimization is performed for a dataset with a fixed repeat interval R. Later, in the experimental simulations model performance is evaluated for other R values, which shows the generalization ability of the model.

We also optimized the baseline LSTM model via Adam algorithm, which is a common gradient-based optimization method (Kingma & Ba, 2014). The baseline LSTM did not have any Hebbian-like mechanisms; its weights were learned during training and then fixed for the test experiments. During training, a new dataset was generated for each backpropagation step, and no separate validation set was required. LSTM was trained until it reached an accuracy of 97%.

### Izhikevich spiking model

Our network consists of 100 spiking neurons, each receiving independent input from one dimension of the input vector. Spiking dynamics was simulated with an Izhikevich neuron model (Izhikevich, 2007; Eq. 7-10). The Izhikevich model can approximate the dynamics of different types of neurons as well as a classical Hodgkin-Huxley model (Hodgkin & Huxley,), but is more efficient, since it has only two differential equations to solve. These equations describe the dynamics of two variables which characterize a neuron state: a membrane potential (voltage) v_i_ and the recovery variable u_i_.

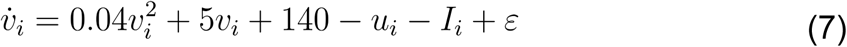

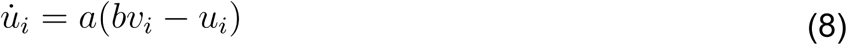

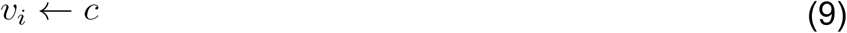

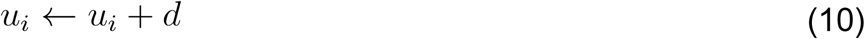

The equations above define how voltage *v_i_* and recovery *u_i_* of neuron *i* evolve over time. They update every time step and are dependent on the neuron total input *I_i_* and voltage noise *ε*. The parameter a is a timescale of recovery. In our model, *a* = 0.02. The parameter *b* describes how sensitive the recovery variable is to the fluctuation of the membrane potential. In our model, *b* = 0.2.

When the voltage reaches the activation threshold 30 mV, we interpret it as a spike event. Note that this threshold rather represents a peak voltage during a spike event. The afterspike neuron dynamics is as follows: the voltage variable is reset according to Eq. 7, and the recovery variable gets updated following Eq. 8. Noise in the model is described in Eq. 11 It is generated randomly at every time step of the dynamical model.

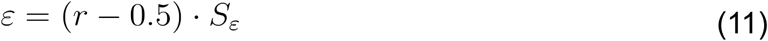

Here, r is a number between 0 and 1 from a uniform distribution. It is adjusted by a voltage noise scaling factor *S_ε_* = 0.3.

The total input to the model at every time step comes from two sources: the external stimulus input and lateral input, i.e. input from the other neurons in the model (Eq. 12).

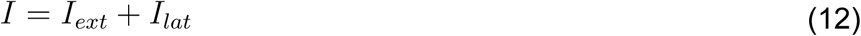

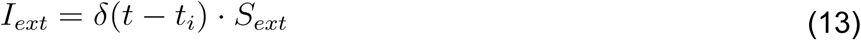

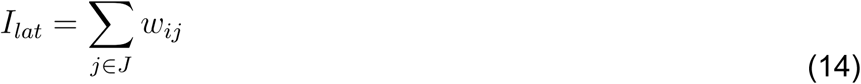

Izhikevich neurons in the model form one-to-one connections with the binary input vector (see Fig. 1). All input weights in the model are similar and equal to a constant external input scaling factor *S_ext_*. Thus, each neuron connected to a non-zero element of the input vector, receives an external input *I_ext_* equal to a scaling factor *S_ext_* at 100 events evenly distributed per 1000 ms via Poisson point process.

In Eq. 14, lateral input *I_lat_*to a neuron i is represented by a summation of weights from the neurons of subset *J*, which are laterally connected to neuron i and emitted a spike at the previous time step. The weights are constantly changing through the ongoing STDP process.

### Measuring model performance

The performance of models in the experiments was measured as prediction accuracy balanced by a proportion of novel stimuli in the dataset. Accuracy in the model depends on the true positive and false positive rate of its predictions (Eq. 15-16, following Tyulmankov et al., 2022).

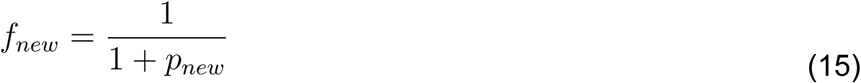

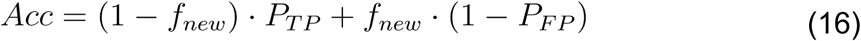

Here, *f_new_* refers to the fraction of novel stimuli in the dataset and depends on *p_new_* **−** the probability to generate a new random stimulus while building a dataset. The accuracy of the model predictions Acc takes into account *P_tp_* and *P_fp_*: the probability of correctly classifying a repeated stimulus as familiar, and incorrectly classifying a randomly generated new stimulus as familiar, respectively.

### Regression analysis

To estimate how well different connectivity features influence the LESH generalizability, we conducted a regression analysis using two methods: Linear regression and Decision Tree regression. Both methods were implemented with the use of the *scikit-learn* package in Python (Pedregosa et al., 2011). Linear regression was used to model the linear relationship, i.e. a weighted sum of the features, between the connectivity features as predictors and the model generalizability on a task of continual familiarity across the range of repeat interval values as a target variable. In contrast, Decision Tree regression splits the predictor feature space into a series of decision-based hierarchical partitions, in order to reduce variance in the target variable within each partition.

We performed 10-fold cross-validation to ensure the robustness of the analysis and prevent overfitting, on a dataset consisting of 360 samples (the data for all optimization repeat intervals and sparseness levels was combined: 3 repeat interval values x 3 sparseness levels x 40 trials). We evaluated two models: with the target generalizability variable estimated from Rsync-based performance, and from rate-based performance.

A set of restrictions was imposed on every model. For Linear regression, we applied the Elastic Net algorithm, which balances L1 and L2 regularization to mitigate overfitting (Zou & Hastie, 2005). The Decision Tree models were constrained to a maximum depth of 5 to maintain interpretability and avoid overfitting. Additionally, the grid search was performed on sets of all connectivity features (Gini index, modularity, transitivity, participation coefficient, betweenness centrality) for both Decision Tree and Linear Regression models, to exclude the features which did not contribute to the model performance, thus selecting the connectivity features important for predicting generalizability. For Linear regression, only betweenness-centrality yielded non-zero regression coefficients, whereas for Decision Tree regression the important features were betweenness-centrality, Gini index and transitivity.

Feature importance was also assessed with automated tools provided by *scikit-learn* (Pedregosa et al., 2011). For Linear regression, the feature importance was computed as a magnitude of the regression coefficients. For Decision Tree regression, it was calculated as a reduction in a node/split impurity, averaged across all nodes in the tree and weighted by the number of samples per node (Eq. 17).

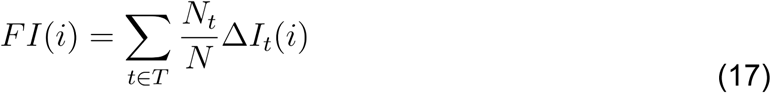

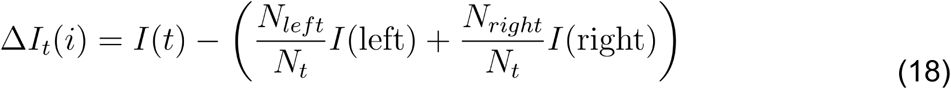

Here, *N* stands for the total number of samples in the dataset, *T* represents a set of nodes in a Decision Tree, *N_t_*is a number of samples per node, and *ΔI_t_(i)* is a reduction in impurity at the node t due to the split based on the value of feature *i*. Eq. 18 defines how the reduction in impurity is calculated: I(t) stands for the impurity of the parent node *t*, *I(left)* and *I(right)* represent the impurity of left and right node children respectively. *N_left_* and *N_right_* stand for the number of samples in left and right children nodes. The node impurity is calculated as a mean-squared error (MSE). It captures how the MSE decreases due to the specific split.

The performance of regression models was evaluated with the root mean square error (RMSE) and the coefficient of determination (R^2^). RMSE is a measure of a predictive error which captures the standard deviation of residuals in a regression, and R^2^ is a goodness-of-fit metric, which quantifies how much variance of the target variable is explained by the model. Thus, a superior model has lower RMSE and higher R^2^ scores (Eq. 19-20).

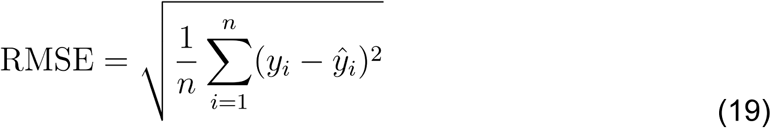

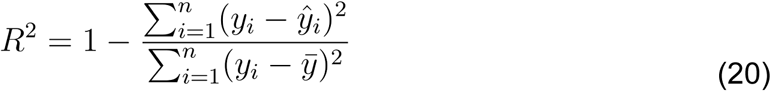

In both formulas, the differences between true and predicted values of *y* are computed, and *n* stands for the number of data points.

## Supplementary

**Table S1.** Fixed model parameters.

| Parameter | Value | Units (where applicable) |
| --- | --- | --- |
| Number of neurons | 100 |  |
| Input firing rate | 100 | Hz |
| External input scaling factor | 21 |  |

**Table S2.** Fixed Izhikevich parameters.

| Parameter | Value | Units (where applicable) |
| --- | --- | --- |
| Simulation time step | 0.5 | ms |
| Initial membrane potential | 30 | mV |
| Initial recovery variable | 30 | mV |
| Izhikevich a (time scale of recovery variable) | 0.01 |  |
| Izhikevich b (sensitivity of recovery variable) | -0.1 |  |
| Izhikevich c (voltage afterspike reset) | -65.0 | mV |
| Izhikevich d (recovery afterspike update) | 12.0 | mV |
| Excitatory reverse potential | 0.0 | mV |
| Voltage noise amount | 0.55 |  |
| Spike detection threshold | 30 | mV |

## Notes

### Competing Interest Statement

The authors have declared no competing interest.

## References

Anderson, B., Mruczek, R. E., Kawasaki, K., & Sheinberg, D. (2008). Effects of familiarity on neural activity in monkey inferior temporal lobe. Cerebral cortex, 18(11), 2540–2552.

Androulidakis, Z., Lulham, A., Bogacz, R., & Brown, M. W. (2008). Computational models can replicate the capacity of human recognition memory. Network: Computation in Neural Systems, 19(3), 161–182.

Antonov, D. I., Sviatov, K. V., & Sukhov, S. (2022). Continuous learning of spiking networks trained with local rules. Neural Networks, 155, 512–522.

Bogacz, R., & Brown, M. W. (2003). Comparison of computational models of familiarity discrimination in the perirhinal cortex. Hippocampus, 13(4), 494–524.

Brown, M. W., Wilson, F. A. W., & Riches, I. P. (1987). Neuronal evidence that inferomedial temporal cortex is more important than hippocampus in certain processes underlying recognition memory. Brain research, 409(1), 158–162.

Bülthoff, I., & Newell, F. N. (2006). The role of familiarity in the recognition of static and dynamic objects. Progress in Brain Research, 154, 315–325.

Cooke, S. F., Komorowski, R. W., Kaplan, E. S., Gavornik, J. P., & Bear, M. F. (2015). Visual recognition memory, manifested as long-term habituation, requires synaptic plasticity in V1. Nature Neuroscience, 18(2), 262–271.

Gerstner, W., & Kistler, W. M. (2002). Mathematical formulations of Hebbian learning. Biological Cybernetics, 87(5), 404–415.

Han, B., Zhao, F., Li, Y., Kong, Q., Li, X., & Zeng, Y. (2024). Similarity-based context aware continual learning for spiking neural networks. Neural Networks, 107037.

Hartigan, J. A., & Hartigan, P. M. (1985). The dip test of unimodality. The Annals of Statistics, 70–84.

Hayden, D. J., Finnie, P. S., Thomazeau, A., Li, A. Y., Cooke, S. F., & Bear, M. F. (2023). Electrophysiological signatures of visual recognition memory across all layers of mouse V1. Journal of Neuroscience, 43(44), 7307–7321.

Hebb, D. (1949). The Organization of Behavior. emphNew York. Wiley.

Hochreiter S, and Schmidhuber J (1997). Long Short-Term Memory. Neural Computation 9, 1735–1780.

Holland, J. H. (1992). Genetic algorithms. Scientific American, 267(1), 66–73.

Hopfield JJ (1982). Neural networks and physical systems with emergent collective computational abilities. PNAS, 79, 2554–2558.

Huang, G., Ramachandran, S., Lee, T. S., & Olson, C. R. (2018). Neural correlate of visual familiarity in macaque area V2. Journal of Neuroscience, 38(42), 8967–8975.

Izhikevich, E. M. (2003). Simple model of spiking neurons. IEEE Transactions on Neural Networks, 14(6), 1569–1572.

Korndörfer, C., Ullner, E., García-Ojalvo, J., & Pipa, G. (2017). Cortical spike synchrony as a measure of input familiarity. Neural Computation, 29(9), 2491–2510.

Kingma, D. P., & Ba, J. (2014). Adam: A method for stochastic optimization. arXiv preprint arXiv:1412.6980.

Lazar, A., Pipa, G., & Triesch, J. (2009). SORN: a self-organizing recurrent neural network. Frontiers in Computational Neuroscience, 3, 800.

Li, T., Tang, M., & Bogacz, R. (2023). Modeling Recognition Memory with Predictive Coding and Hopfield Networks. In Associative Memory & Hopfield Networks.

Liang, F., Li, H., Chou, X. L., Zhou, M., Zhang, N. K., Xiao, Z., … & Zhang, L. I. (2019). Sparse representation in awake auditory cortex: cell-type dependence, synaptic mechanisms, developmental emergence, and modulation. Cerebral Cortex, 29(9), 3796–3812.

Lindsey, J., & Litwin-Kumar, A. (2020). Learning to learn with feedback and local plasticity. Advances in Neural Information Processing Systems, 33, 21213–21223.

Meyer, T., & Rust, N. C. (2018). Single-exposure visual memory judgments are reflected in inferotemporal cortex. elife, 7, e32259.

Miller, E. K., Li, L., & Desimone, R. (1991). A neural mechanism for working and recognition memory in inferior temporal cortex. Science, 254(5036), 1377–1379.

Norman, K. A., & O’Reilly, R. C. (2003). Modeling hippocampal and neocortical contributions to recognition memory: a complementary-learning-systems approach. Psychological Review, 110(4), 611.

Olshausen, B. A., & Field, D. J. (2004). Sparse coding of sensory inputs. Current opinion in neurobiology, 14(4), 481–487.

Pedregosa, F., Varoquaux, G., Gramfort, A., Michel, V., Thirion, B., Grisel, O., … & Duchesnay, É. (2011). Scikit-learn: Machine learning in Python. The Journal of Machine Learning Research, 12, 2825–2830.

Pipa, G., Wheeler, D. W., Singer, W., & Nikolić, D. (2008). NeuroXidence: reliable and efficient analysis of an excess or deficiency of joint-spike events. Journal of Computational Neuroscience, 25, 64–88.

Rainer, G., & Miller, E. K. (2000). Effects of visual experience on the representation of objects in the prefrontal cortex. Neuron, 27(1), 179–189.

Ringo, J. L. (1996). Stimulus specific adaptation in inferior temporal and medial temporal cortex of the monkey. Behavioural brain research, 76(1-2), 191–197.

Rubinov, M., & Sporns, O. (2010). Complex network measures of brain connectivity: uses and interpretations. Neuroimage, 52(3), 1059–1069.

Shen, J., Ni, W., Xu, Q., & Tang, H. (2024). Efficient spiking neural networks with sparse selective activation for continual learning. In Proceedings of the AAAI Conference on Artificial Intelligence (Vol. 38, No. 1, pp. 611–619).

Terrell, G. R., & Scott, D. W. (1992). Variable kernel density estimation. The Annals of Statistics, 1236–1265.

Toutounji, H., & Pipa, G. (2014). Spatiotemporal computations of an excitable and plastic brain: neuronal plasticity leads to noise-robust and noise-constructive computations. PLoS Computational Biology, 10(3), e1003512.

Tyulmankov, D., Yang, G. R., & Abbott, L. F. (2022). Meta-learning synaptic plasticity and memory addressing for continual familiarity detection. Neuron, 110(3), 544–557.

Xiang, J. Z., & Brown, M. W. (1998). Differential neuronal encoding of novelty, familiarity and recency in regions of the anterior temporal lobe. Neuropharmacology, 37(4-5), 657–676.

Yonelinas, A. P. (2001). Components of episodic memory: the contribution of recollection and familiarity. Philosophical Transactions of the Royal Society of London. Series B: Biological Sciences, 356(1413), 1363–1374.

Yonelinas, A. P. (2002). The nature of recollection and familiarity: A review of 30 years of research. Journal of memory and language, 46(3), 441–517.

Yonelinas, A. P., Kroll, N. E. A., Dobbins, I. G., & Soltani, M. (1999). Recognition memory for faces: When familiarity supports associative recognition judgments. Psychonomic Bulletin & Review, 6(4), 654–661.

Yoshida, T., & Ohki, K. (2020). Natural images are reliably represented by sparse and variable populations of neurons in visual cortex. Nature communications, 11(1), 872.

Zemliak, V., Mayer, J., Nieters, P., & Pipa, G. (2024). Spike synchrony as a measure of Gestalt structure. Scientific Reports, 14(1), 5910.

Zou, H., & Hastie, T. (2005). Regularization and variable selection via the elastic net. Journal of the Royal Statistical Society Series B: Statistical Methodology, 67(2), 301–320

